# Striatal dopamine synthesis and cognitive flexibility differ between hormonal contraceptive users and non-users

**DOI:** 10.1101/2022.10.20.513082

**Authors:** Caitlin M. Taylor, Daniella J. Furman, Anne S. Berry, Robert L. White, William J. Jagust, Mark D’Esposito, Emily G. Jacobs

## Abstract

In rodents and nonhuman primates, sex hormones are powerful modulators of dopamine neurotransmission. Yet little is known about hormonal regulation of the dopamine system in the human brain. Using Positron Emission Tomography (PET), we address this gap by comparing hormonal contraceptive users and non-users across multiple aspects of dopamine function: dopamine synthesis capacity via the PET radioligand 6-[^18^F]fluoro-m-tyrosine ([^18^F]FMT), baseline D2/3 receptor binding potential using [^11^C]raclopride, and dopamine release using methylphenidate-paired [^11^C]raclopride. Participants consisted of 36 healthy women (n=21 naturally cycling; n=15 hormonal contraceptive users), and men (n=20) as a comparison group. A behavioral index of cognitive flexibility was assessed prior to PET imaging. Hormonal contraceptive users exhibited greater dopamine synthesis capacity than naturally cycling participants, particularly in dorsal caudate, and greater cognitive flexibility. Further, across individuals the magnitude of striatal DA synthesis capacity was associated with cognitive flexibility. No group differences were observed in D2/3 receptor binding or dopamine release. Analyses by sex alone may obscure underlying differences in DA synthesis tied to women’s hormone status. Hormonal contraception (in the form of pill, shot, implant, ring or IUD) is used by ~400 million women worldwide, yet few studies have examined whether chronic hormonal manipulations impact basic properties of the dopamine system. Findings from this study begin to address this critical gap in women’s health.

---

Sex hormones are powerful neuromodulators of learning and memory (1). Accumulating evidence suggests that sex hormones’ influence extends to the regulation of dopamine (DA) (2–5), itself a neuromodulator of higher order cognitive functions (6–8). In rodents and nonhuman primates, 17β-estradiol (E2) and progesterone (P) modulate DA synthesis and release, alter DA-D2 receptor availability, and modify the basal firing rate of dopaminergic neurons (9–15). For example, E2 administration produces a dose-dependent increase in striatal DA (11) and modulates goal-directed behavior (16) in rodents. Progesterone has a bimodal effect on striatal DA concentration, with increases in DA in the first 12 hours after P perfusion, and inhibitory effects 24h post-infusion. Further, surgical removal of the ovaries reduces tyrosine hydroxylase–immunoreactive neurons in the substantia nigra (17) and prefrontal cortex (18). Estrogen receptors are localized to regions that receive major projections from midbrain DA neurons, including prefrontal cortex (PFC), dorsal striatum, and the nucleus accumbens (19). Despite the substantial literature supporting sex hormones’ role in DA neuromodulation in rodents and nonhuman primates, little is known about hormonal regulation of the dopamine system in the human brain.

Indirect evidence in humans suggests that estradiol modulates dopamine-dependent cognitive function and prefrontal cortex activity (20–22, 22, 23). For example, Jacobs and D’Esposito found evidence that estradiol regulates PFC activity and working memory performance, and the direction of the effect depends on an individual’s basal PFC dopamine tone (indexed by catechol-O-methyltransferase activity) (20). Additional evidence suggests that menstrual cycle phase influences dopamine-dependent motor and cognitive functions, including response time on tests of manual coordination, working memory and cognitive flexibility (24,25), and immediate reward selection bias (26).

Molecular PET imaging provides a more direct assessment of dopaminergic activity *in vivo* in the human brain. Findings of sex differences in DA synthesis capacity (27), DA release (28–30), and DA transporter density (31,32) again suggest a role for sex steroid hormones in modulating aspects of DA functioning. Additional evidence comes from PET studies of women in different phases of the menstrual cycle or during the menopausal transition. Wong et al. (33) observed fluctuations in DA-D2 receptor density across the menstrual cycle in healthy premenopausal women, and Pohjalainen et al. (34) observed greater variability in DA-D2 receptor density in premenopausal versus postmenopausal women, with the suggestion that greater variability was attributable to hormonal fluctuations across the menstrual cycle. Evidence is mixed, however, with some studies reporting no significant associations between DA signaling and menstrual cycle phase or serum estradiol concentrations (35–37).

An underexplored population for studying hormonal influences on DA function is women using hormonal contraception. Hormonal contraception (HC; in the form of pill, shot, implant, ring or IUD) is used by ~400 million women worldwide (38), yet few studies have examined whether chronic hormone manipulations affect basic properties of the dopamine system. In the present study, we probed the impact of hormonal contraception on multiple properties of the DA system using molecular PET imaging techniques, offering new insights into the relationship between sex hormones and DA neurotransmission in the human brain. The study consisted of young, healthy women and men, and paired pharmacological manipulation of the DA system with PET imaging to assess synthesis capacity (radioligand [^18^F]fluoro-m-tyrosine), D2 receptor availability (radioligand [^11^C]raclopride) and DA release (radioligand [^11^C]raclopride paired with methylphenidate). This provides a unique opportunity to characterize differences in DA synthesis capacity, basal DA receptor occupancy, and stimulated DA release in a single cohort. Next, we investigated sex differences in DA neurotransmission. Finally, we examined whether differences in DA neurotransmission were associated with DA-dependent cognition, using a behavioral assessment of cognitive flexibility (39,40).

## Methods

### Participants

Participants consisted of 57 adults (mean age = 21.16, SD =2.37, range: 18–28 years), including 37 women and 20 men (n = 28 Asian, 10 Hispanic or Latino, 9 White (not Hispanic or Latino), 2 Black or African-American, 3 more than one race or ethnicity, 2 other, and 3 declined to state). Participants underwent PET and MRI imaging as part of a parent study on dopaminergic mechanisms of cognitive control (e.g., see (40)). PET data from this sample have previously been described in (41). This study was approved by Institutional Review Boards at the University of California, Berkeley and Lawrence Berkeley National Laboratory. Participants met the following eligibility criteria: (1) 18–30 years old, (2) right-handed, (3) current weight of at least 100 pounds, (4) able to read and speak English fluently, (5) nondrinker or light drinker (women: <7 alcoholic drinks/week; men: <8 alcoholic drinks/week), (6) no recent history of substance abuse, (7) no history of neurological or psychiatric disorder as confirmed by clinician interview, (8) no current psychoactive medication or street drug use, (9) not pregnant, and (10) no contraindications to MRI. Most participants completed three PET scans over the course of two separate sessions: [18F]FMT, and [^11^C]raclopride + placebo and [^11^C]raclopride + methylphenidate on the same day; the exceptions were one participant (a naturally-cycling woman) who did not complete the FMT scan due to technical issues, one participant (a naturally-cycling woman) who did not produce reliable Raclopride scan data due to technical issues, and two participants (hormonal contraceptive users) who did not complete Raclopride scans.

#### FMTsample

Women were categorized based on hormone status: naturally cycling (NC, no current reported use of hormonal contraception; n = 21, avg. age = 22.67 years, SD = 2.77) and current users of hormonal contraception (HC, n = 15, avg. age = 20.43 years, SD = 1.91). Types of hormonal contraception used included: combined oral contraception (OC, n = 10), vaginal ring (n = 1), implant (n = 2), injection (n = 1), and hormonal intrauterine device (IUD, n = 1).

#### RAC sample

RAC data from one NC participant did not pass quality control and two HC users (combined OC) did not have RAC sessions, yielding a final sample of 21 NC women (avg age = 20.67, SD = 1.91) and 13 HC users (avg age = 22.69, SD = 2.81).

In our secondary analyses, participants were grouped by self-reported sex (male, n = 20; female, n = 37), and hormone status (male, NC, HC). Men and women did not differ significantly in age or BMI, however HC users were older than males (*p* = .03, *d* = 0.85) and NC participants (*p* = .01, *d* = 0.94) by 25 months on average (Table 1).

**Table 1.** Participant demographics by sex and hormone status

|  | Age | BMI |
| --- | --- | --- |
| **Men** (n= 20) | 20.7 ± 2.1 | 23.8 ± 5.3 |
| **Women** (n = 37) | 21.4 ± 2.5 | 23.7 ± 4.2 |
| **Naturally Cycling**<br>(NC, n = 22) | 20.6 ± 2.0 | 23.0 ± 3.9 |
| **Hormonal Contraception**<br>(HC, n = 15) | 22.7 ± 2.8 | 24.7 ± 4.6 |
| **NC vs HC**<br>cohen's d (Welch's *p*) | 0.94 (.01<sup>1</sup>) | n.s. |
| **Men vs Women** | n.s. | n.s. |
| **Men vs NC vs HC**<br>Kruskal–Wallis *p* | .03 | n.s. |

<sup>1</sup>Indicates significance with Bonferroni correction ( $p < .0167$ )
Types of hormonal contraception used (n): Combined OC (10), Vaginal ring (1), Implant (2), Injection (1), Hormonal IUD (1)

### Structural MRI Scan

Images were acquired using a Siemens 3 T Trio Tim scanner with a 12-channel coil. Each participant was scanned on 3 occasions using a high-resolution T1-weighted magnetization prepared rapid gradient echo (MPRAGE) whole brain scan (TR=2,300 ms; TE=2.98 ms; FA=9°; matrix=240 × 256; FOV=256; sagittal plane; voxel size=1 × 1 × 1 mm; 160 slices). The three MPRAGE scans were aligned, averaged, and segmented using FreeSurfer version 5.1 (http://surfer.nmr.mgh.harvard.edu/) and the averaged template was used for coregistration with the PET data.

### [^18^F]FMT PET Data Acquisition

Participants underwent an [^18^F]FMT PET scan to measure dopamine synthesis capacity. Detailed methods are provided in (39). PET data were acquired using a Siemens Biograph Truepoint 6 PET/CT scanner (Siemens Medical Systems, Erlangen, Germany) ~1 hour after participants ingested 2.5 mg/kg of carbidopa to minimize the peripheral decarboxylation of [^18^F]FMT. After a short CT scan, participants were injected with approximately 2.5 mCi of [^18^F]FMT as a bolus in an antecubital vein (M ±SD; specific activity = 947.30 ± 140.26 mCi/mmol; dose= 2.43 ±0.06 mCi). Dynamic acquisition frames were obtained over 90 min in 3D mode (25 frames total: 5 × 1, 3 × 2, 3 × 3, 14 × 5 min). Data were reconstructed using an ordered subset expectation maximization algorithm with weighted attenuation, corrected for scatter, and smoothed with a 4mm full width at half maximum (FWHM) kernel.

### [11C]Raclopride PET Data Acquisition

Participants received two [^11^C]raclopride PET scans an average of 21.65 days before or after the [^18^F]FMT scan (median = 7 days) to measure D2/3 receptor occupancy and dopamine release. To measure baseline D2/3 receptor occupancy, participants ingested a placebo pill approximately 1 hour before [^11^C]raclopride scan 1. The placebo scan was always performed first. To measure dopamine release, participants ingested 30 mg (M ± SD mg/kg: 0.46 ± 0.08) of methylphenidate ~ 1 hour before [^11^C] raclopride scan 2. Endogenous DA release was measured as the percent signal change (PSC) in non-displaceable binding potential (BPND) from [^11^C]raclopride scan 1 to [^11^C]raclopride scan 2 ((placebo [^11^C]raclopride - methylphenidate [^11^C]raclopride)/placebo [^11^C]raclopride). Scans were conducted on the same day, 2 hours apart and participants were blind to whether placebo or methylphenidate was administered. For both [^11^C]raclopride scan 1 and [^11^C] raclopride scan 2, after a short CT scan, participants were injected with approximately 10 mCi of [^11^C]raclopride as a bolus in an antecubital vein. Dynamic acquisition frames were obtained over 60 min in 3D mode (19 frames total: 5 × 1, 3 × 2, 3 × 3, 8 × 5). Reconstruction was performed as described above.

### PET Data Analysis

PET data were preprocessed using SPM8 software (Friston et al, 2007). To correct for motion between frames, images were realigned to the middle frame. The first five images were summed prior to realignment to improve realignment accuracy, as these early images have relatively low signal contrast. Structural images were coregistered to PET images using the mean image of frames corresponding to the first 20 min of acquisition as a target. The mean image for the first 20 min was used rather than the mean image for the whole scan time because it provides a greater range in image contrast outside of striatum thus making it a better target for coregistration.

#### [^18^F]FMT

Graphical analysis for irreversible tracer binding was performed using Patlak plotting (42,43) implemented using inhouse software and Matlab version 8.2 (The MathWorks, Natick, MA). Without measurement of the arterial input function [^18^F]FMT PET analysis used reference region models. Cerebellar gray matter was used as the reference region because this region shows very little tracer uptake, and has an extremely low density of DA receptors and metabolites relative to striatum (44–47). The most anterior ¼ of cerebellar gray was removed from the reference region to limit contamination of signal from the substantia nigra and ventral tegmental area. K_i_ images were generated from PET frames corresponding to 25 to 90min (48,49), which represent the amount of tracer accumulated in the brain relative to the reference region.

#### [^11^C]Raclopride

For [^11^C]raclopride PET, reversible tracer binding was quantified using simplified reference tissue model analysis (SRTM; (50)). Specifically, a basis function version of the SRTM was applied as previously described (51) with posterior cerebellar gray matter used as the reference region. The SRTM analysis was performed using inhouse software provided by Dr Roger Gunn and Matlab version 8.2. SRTM analysis was used to determine BPND, which can be defined as: BP_ND_= f_ND_B_avail_/K_D_ where B_avail_ is the concentration of D2/3 receptors, K_D_ is the inverse of the affinity of the radiotracer for D2/3 receptors, and f_ND_ is the free fraction of the ligand in the nondisplaceable tissue compartment (52,53).

### Regions of Interest

An ROI approach was used to test relationships between hormonal status and PET measures of dopaminergic function in striatal subregions. Striatal subregions were manually drawn in native space on each participant’s averaged MPRAGE MRI scan using Mango software. The dorsal caudate, dorsal putamen, and ventral striatum were segmented as described in (54). Inter-rater reliability was high for manually drawn striatal subregions (see (39)).

As we did not hypothesize an effect of hemisphere, ROI values for our three ROIs (dorsal caudate, dorsal putamen, and ventral striatum) were analyzed as voxel-weighted averages of left and right hemisphere PET values as follows:

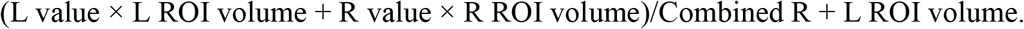

All analyses on striatal values were conducted on partial volume corrected ROIs (PVC; (55)). PVC was performed in native space (non-normalized data) and corrects for between-subject differences in the inclusion of white matter and CSF in the measured volumes. To apply the PVC in native space, we used FreeSurfer-generated ROIs for gray matter cortical and subcortical regions, white matter, and cerebral spinal fluid. All statistical analyses were conducted using R (version 1.2.5001)

### Cognitive Paradigm

The task was an adaptation of the task-switching paradigm developed by Armbruster, Ueltzhöffer, Basten, and Fiebach (56) and is described in detail in (40). Briefly, on each trial, participants were required to respond quickly to digits between 1 and 9 (excluding 5) that appeared in different shades of gray against a black background. On 82% of trials, a single digit appeared above a central fixation. For these “ongoing task” trials, participants performed an operation (odd/even or low/high decisions) on the digit and responded by pressing the index finger of either their left or right hand. On the remaining 18% of trials, two digits appeared on the screen simultaneously, one above and one below the fixation cross. The relative brightness of the upper and lower digits varied and encoded a task cue. When the upper digit was brighter (6% of trials), participants were instructed to ignore the lower digit and continue to apply the ongoing task rule to the upper digit (“distractor trials”). When the lower digit was brighter (6% of trials), participants were signaled to switch attention to the lower and to apply the alternate task rule to it (“switch trials”). On the final third of these trials (6% of total trials), the difference in brightness between the upper and lower digits was reduced (“ambiguous trials”). Ambiguous trials were not considered in our analyses. Participants performed a total of 990 trials distributed across three blocks with brief interposed breaks. Cognitive testing occurred prior to PET imaging. *Distractor cost* was calculated as the difference between performance accuracy on “distractor” trials and “ongoing” trials, and *switch cost* was calculated as the difference between performance accuracy on “switch” versus “ongoing” trials. One NC participant did not undergo cognitive testing, resulting in a final sample of 20 NC women (avg age = 20.67, SD = 1.91) and 15 HC users (avg age = 22.69, SD = 2.81)

### Statistical Analysis

#### Impact of hormone status on DA neurotransmission

Since hormonal contraceptive (HC) users were older than naturally cycling (NC) participants, to compare markers of dopaminergic signaling between HC and NC groups, we conducted 2 × 4 ANCOVA (hormone group × bilateral region of interest, controlling for the effects of age) for measures of interest (FMT K_i_, [^11^C]raclopride BP_ND_ and percent signal change (PSC) in [^11^C]raclopride BP). We investigated significant main effects with post-hoc one-way ANCOVAs to determine which regions were driving the effect, controlling for the effects of age. Statistically significant findings that survived Bonferroni correction for multiple comparisons are noted (*p*_Bf_ .05/3 regions = .0167). Partial effect sizes (η^2^) are reported for statistically significant findings.

Welch’s *t*-tests were used to compare distractor costs and switch costs between our comparison groups. One NC participant with unusable task data was omitted from these analyses. Finally, as a follow-up to observed differences between NC and HC women, switch costs were correlated with [^18^F]FMT K_i_ PVC striatal values to evaluate a relationship between performance and DA synthesis.

#### Sex differences in DA neurotransmission

To compare aspects of DA signaling by sex and hormone status, we conducted 2 × 3 mixed ANCOVA (group × bilateral region of interest, controlling for age) for measures of interest (FMT K_i_, [^11^C]raclopride BP and percent signal change (PSC) in [^11^C]raclopride BP). Welch’s *t*-tests were conducted to compare distractor costs and switch costs by sex.

Finally, to determine whether differences in hormonal status within women influenced the detection of sex differences, we conducted 3 × 3 mixed ANCOVA (group × bilateral region of interest, controlling for age) for each measure of interest (FMT K_i_, [^11^C]raclopride BP and percent signal change (PSC) in [^11^C]raclopride BP). Significant main effects were investigated using post-hoc one-way ANCOVAs, again to control for the effects of age.

## Results

### DA neurotransmission differs with hormonal contraceptive use

#### Striatal [^18^F]FMT K_i_

[^18^F]FMT PET data was obtained to assess DA synthesis capacity in the striatum. ANCOVA revealed significant main effects of age (F(1,33) = 4.844, *p* = .035, η^2^ = 0.13), hormone status (F(1,33) = 7.753, *p* = .009, η^2^ = 0.19, **Fig. 1**) and region (F(2,68) = 207.859, *p* < .00001, η^2^ = 0.86). Regional effects were expected as previously reported (41). There was no significant interaction between hormone status and region. Results from post-hoc one-way ANCOVAs indicate that hormonal contraceptive users exhibited greater FMT K_i_ values compared to naturally cycling participants, with the largest effect in dorsal caudate (F(1,33) = 9.611, *p_Bf_*=.004). K_i_ values differed marginally between hormonal contraceptive users and naturally cycling participants in dorsal putamen (F(1,33) = 3.966, *p* = .055) and ventral striatum (F(1,33) = 3.754, *p* = .061) (**Table 2**).

**Figure 1.**
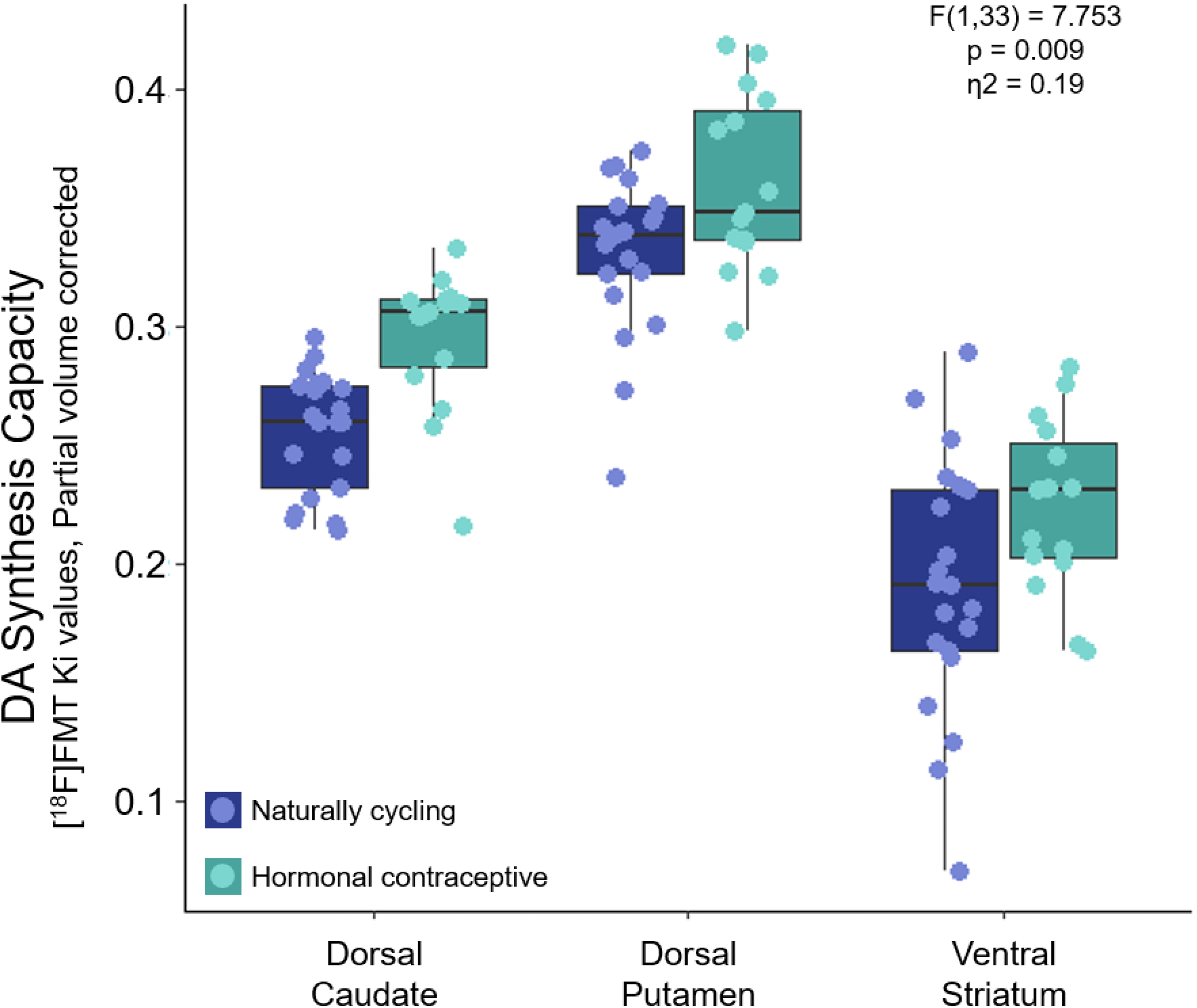
Effect of hormone status on DA synthesis capacity. [^18^F]FMT Ki values in naturally cycling females and hormonal contraceptive users by striatal region of interest. Striatal DA synthesis capacity was greater in hormonal contraceptive users relative to naturally cycling women, with the most pronounced effects observed in dorsal caudate.

**Table 2.**
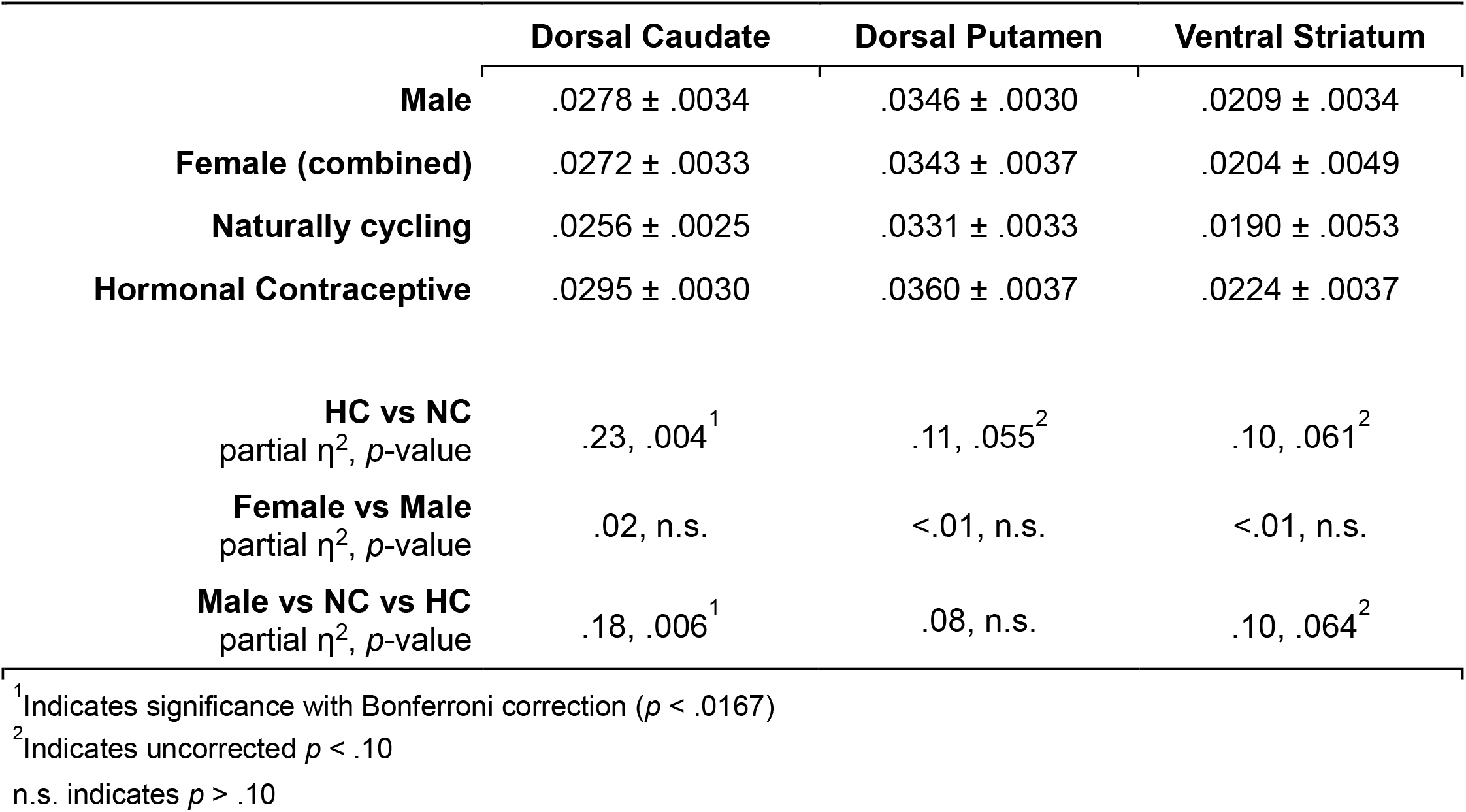
Dopamine synthesis capacity ([^18^F]FMT Ki Values) by group and striatal region of interest

|  | Dorsal Caudate | Dorsal Putamen | Ventral Striatum |
| --- | --- | --- | --- |
| **Male** | .0278 ± .0034 | .0346 ± .0030 | .0209 ± .0034 |
| **Female (combined)** | .0272 ± .0033 | .0343 ± .0037 | .0204 ± .0049 |
| **Naturally cycling** | .0256 ± .0025 | .0331 ± .0033 | .0190 ± .0053 |
| **Hormonal Contraceptive** | .0295 ± .0030 | .0360 ± .0037 | .0224 ± .0037 |
| **HC vs NC**<br>partial $\eta^2$, $p$-value | .23, .004<sup>1</sup> | .11, .055<sup>2</sup> | .10, .061<sup>2</sup> |
| **Female vs Male**<br>partial $\eta^2$, $p$-value | .02, n.s. | <.01, n.s. | <.01, n.s. |
| **Male vs NC vs HC**<br>partial $\eta^2$, $p$-value | .18, .006<sup>1</sup> | .08, n.s. | .10, .064<sup>2</sup> |

<sup>1</sup>Indicates significance with Bonferroni correction ( $p < .0167$ )
<sup>2</sup>Indicates uncorrected $p < .10$
n.s. indicates $p > .10$

#### Striatal [^11^C]Raclopride BP

[^11^C]Raclopride PET data was obtained to measure D2/3 receptor binding potential. [^11^C]Raclopride BP differed significantly by region (F(2,64) = 389.281,*p* < .0001, η^2^ = 0.92). Regional effects were expected as previously reported (41). There was no significant main effect of age (F(1,31) = 3.795, *p* = .061) or hormone status (F(1,31) = 0.09, *p* = .76) on [^11^C]raclopride BP_ND_ values, nor was there an interaction between hormone status and region (F(2,64) = 0.815, *p* = .447) (see Supplemental Table 1 for values).

#### Percent Signal Change in Striatal [^11^C]Raclopride BP

Methylphenidate-paired [11C]raclopride PET data was acquired to measure DA release.

[^11^C]Raclopride BP PSC values differed significantly by region (F(2,64) = 389.281,*p* < .0001, η^2^ = 0.92). Again, regional effects were expected as previously reported (41). There were no significant effects of age (F(1,31) = 3.795, *p* = .061) or hormone status (F(1,31) = 0.092, *p* = .76) on [^11^C]raclopride BP PSC values, nor was there an interaction between status and region (F(2,64) = 0.815, *p* = .45) (see supplemental Table 1 for values).

### DA neurotransmission does not differ by sex

#### Striatal [^18^F]FMT K_i_

We observed a main effect of region on FMT values (F(2,108) = 358.424,*p* < .0001, η^2^ = 0.87), no main effect of sex (F(1,53) = 0.415, *p* = .52), and no interaction between sex and region (F(2,108) = .032, *p* = .97).

#### Striatal [^11^C]Raclopride BP

We observed a main effect of region on [^11^C]raclopride BP values (F(2,104) = 479.362, *p* < .0001, η^2^ = 0.90), but no main effect of sex (F(1,52) = 0.084, *p* = .77), and no interaction between sex and region (F(2,104) = 1.453, *p* = .24).

#### Striatal [^11^C]Raclopride BP Percent Signal Change

Again, we observed a main effect of region on percent signal change in [^11^C]raclopride BP values (F(2,104) = 5.383, *p* = .006, η^2^ = 0.09, but no main effect of sex (F(1,51) = 0.089, *p* = .77), and no interaction between sex and region (F(2,104) = 1.488, *p* = .23).

### Differences in DA neurotransmission by sex and hormone status

#### Striatal [^18^F]FMT K_i_

Despite differences in striatal DA synthesis capacity within women based on hormone status, men did not differ appreciably from women in either hormone group. ANCOVA revealed significant main effects of group (F(2,52 = 5.058, *p* = .010, η^2^ = 0.16) and region (F(2,106) = 116.5, *p* < .00001, η^2^ = 0.60) (**Fig. 2)**. There was no significant effect of age (F(1,52) = 1.444, *p* = .235) and no interaction between group and region (F(4,106) = 0.166, *p* = .96). Post-hoc Tukey’s HSD test confirmed that the main effect of hormone status was driven by previously reported significant differences between naturally cycling and hormonal contraceptive groups (*p* = .004), with no differences between males vs. HC (*p* = .20) or vs. NC (*p* = .12).

**Figure 2.**
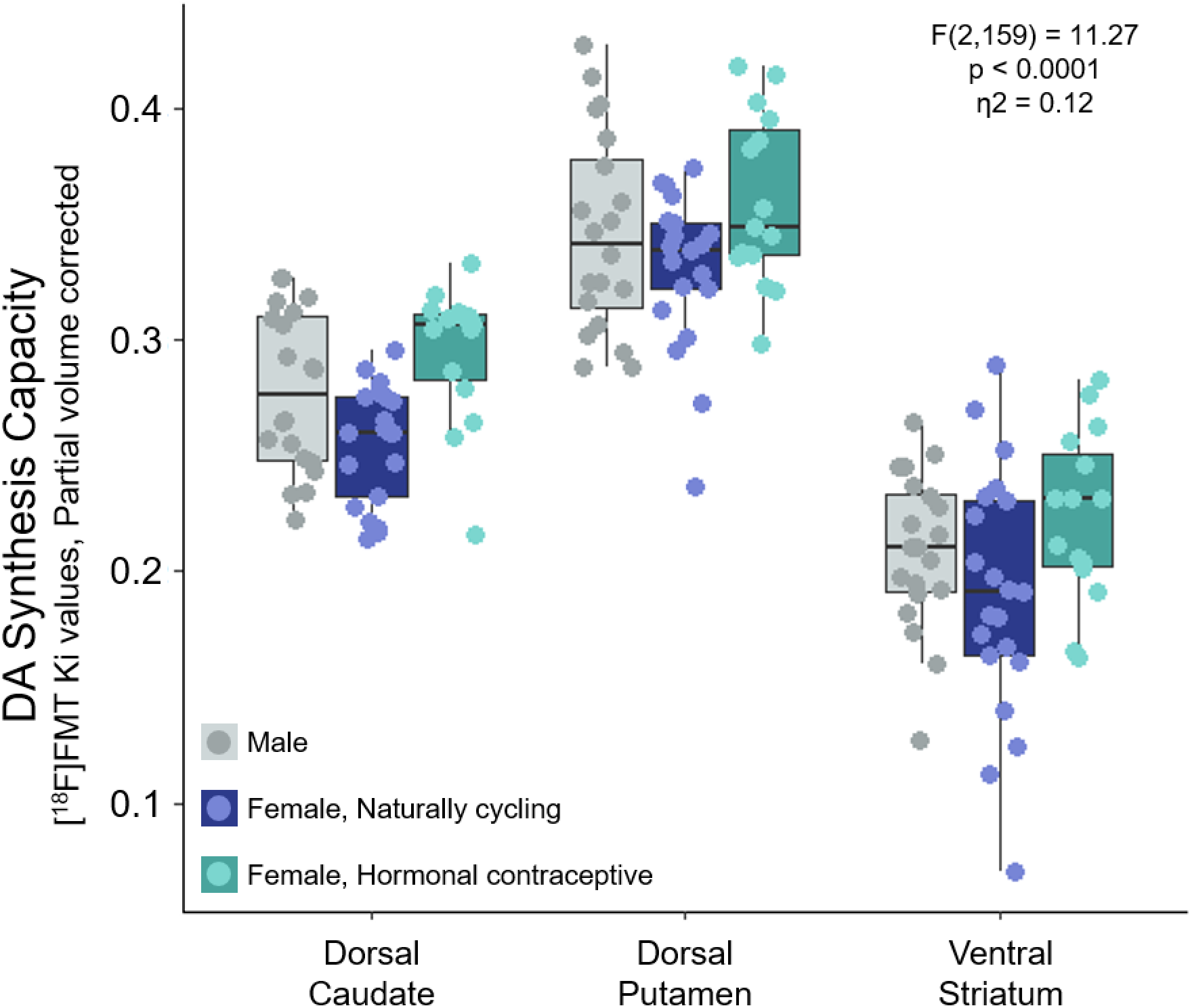
No evidence for sex differences in DA synthesis. [^18^F]FMT Ki values in males, naturally cycling females and hormonal contraceptive users by striatal region of interest. There were no significant differences between males and females (as a whole or by hormone status). As before, striatal DA synthesis capacity was greater in hormonal contraceptive users relative to naturally cycling women.

#### Striatal [^11^C]Raclopride BP

We identified a significant effect of region (F(2,102) = 476.183, *p* < .0001, η^2^ = 0.90), however there was no significant main effect of age (F(1,50) = 1.330, *p* = .25) or hormone status (F(2,50) = 0.044, *p* =.96), nor an interaction between the two factors (F(4,102) = 1.049, *p* = .39).

#### Striatal [^11^C]Raclopride BP PSC

There was a significant effect of region (F(2,102) = 5.284, *p* = .007, η^2^ = 0.09), no significant effect of age (F(1,50) = 0.400, *p* = .53) or hormone status (F(2,50) = 0.081, *p* = .92), and no significant interaction between the two (F(4,012) = 0.750, *p* = .56).

### Individual differences in DA transmission are tied to differences in cognitive flexibility

#### Naturally Cycling vs Hormonal Contraceptive Users

There was no statistically significant difference in *distractor cost* between hormonal contraceptive users and naturally cycling participants (t(31.9) = 0.093, *p* = .926; **Fig. 3A**). However, hormonal contraceptive users exhibited significantly reduced *switch cost* compared to naturally cycling participants (*t*(31.0) = −2.256, *p* = .031; *d* = −0.74; age-adjusted) (**Fig. 3B**). Across female participants, switch cost was inversely correlated with [^18^F]FMT K_i_ values in the dorsal caudate (Pearson’s r(33) = −0.41, *p* = .016) and ventral striatum (r(33) = −0.34, *p* = .042), but not in the dorsal putamen (r(33) = −0.29, *p* = .089). Only the effect in the dorsal caudate was statistically significant after correcting for multiple comparisons (**Fig. 4)**. By contrast, there were no significant correlations between [^18^F]FMT K_i_ values and distractor cost in any ROI (all ps > .6). There were no significant correlations among males between [18F]FMT K_i_ values and switch or distractor costs in any ROI (p > .2 for all).

**Figure 3.**
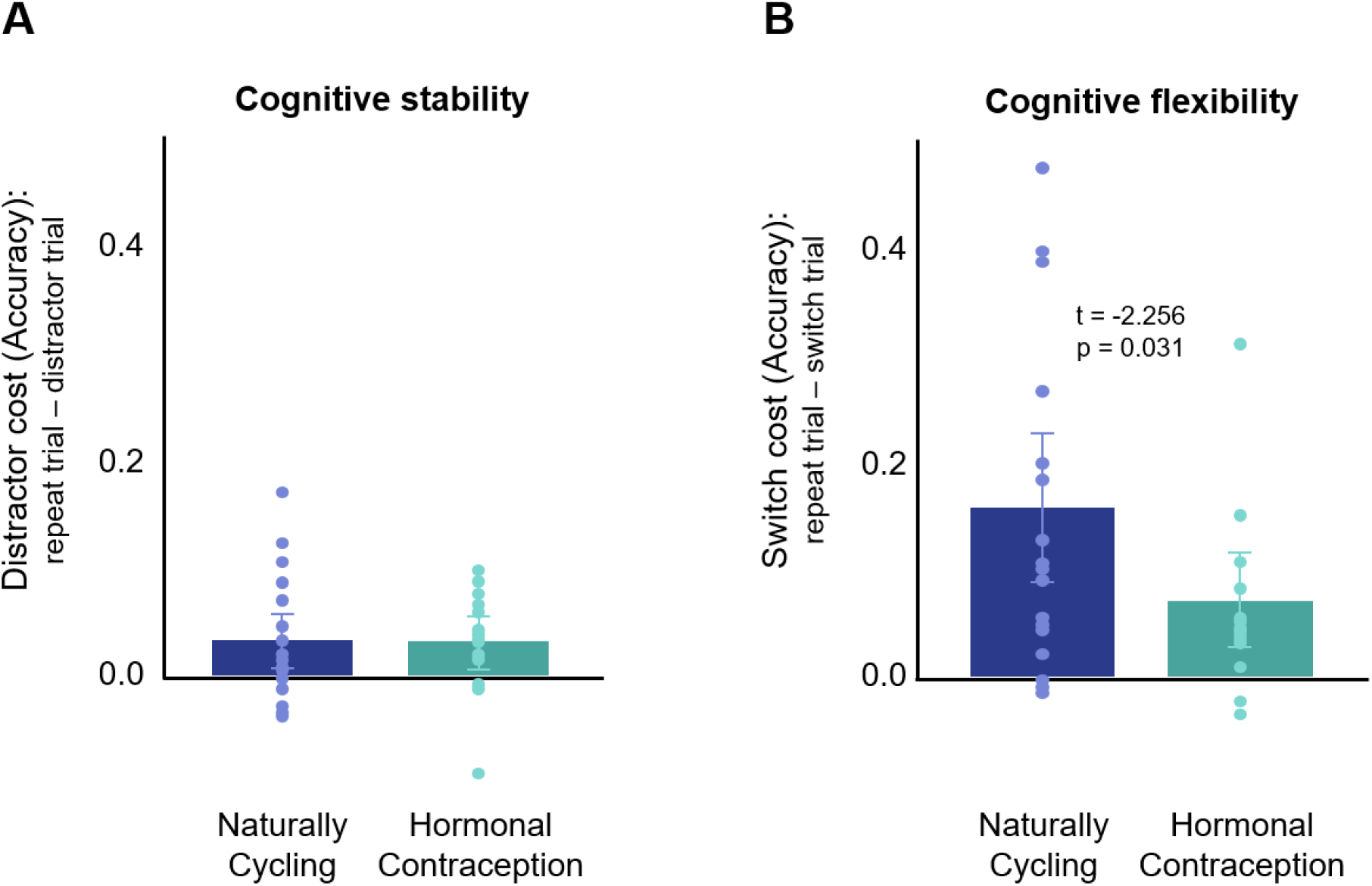
Cognitive flexibility differs between naturally cycling and hormonal contraceptive groups. Performance on a task switching paradigm reveals no difference in cognitive stability between groups, indicated by no difference in distractor costs on distractor/ongoing trials. In contrast, hormonal contraceptive users exhibited greater cognitive flexibility compared to naturally cycling participants, indicated by a smaller performance cost on task-switching trials

**Figure 4.**
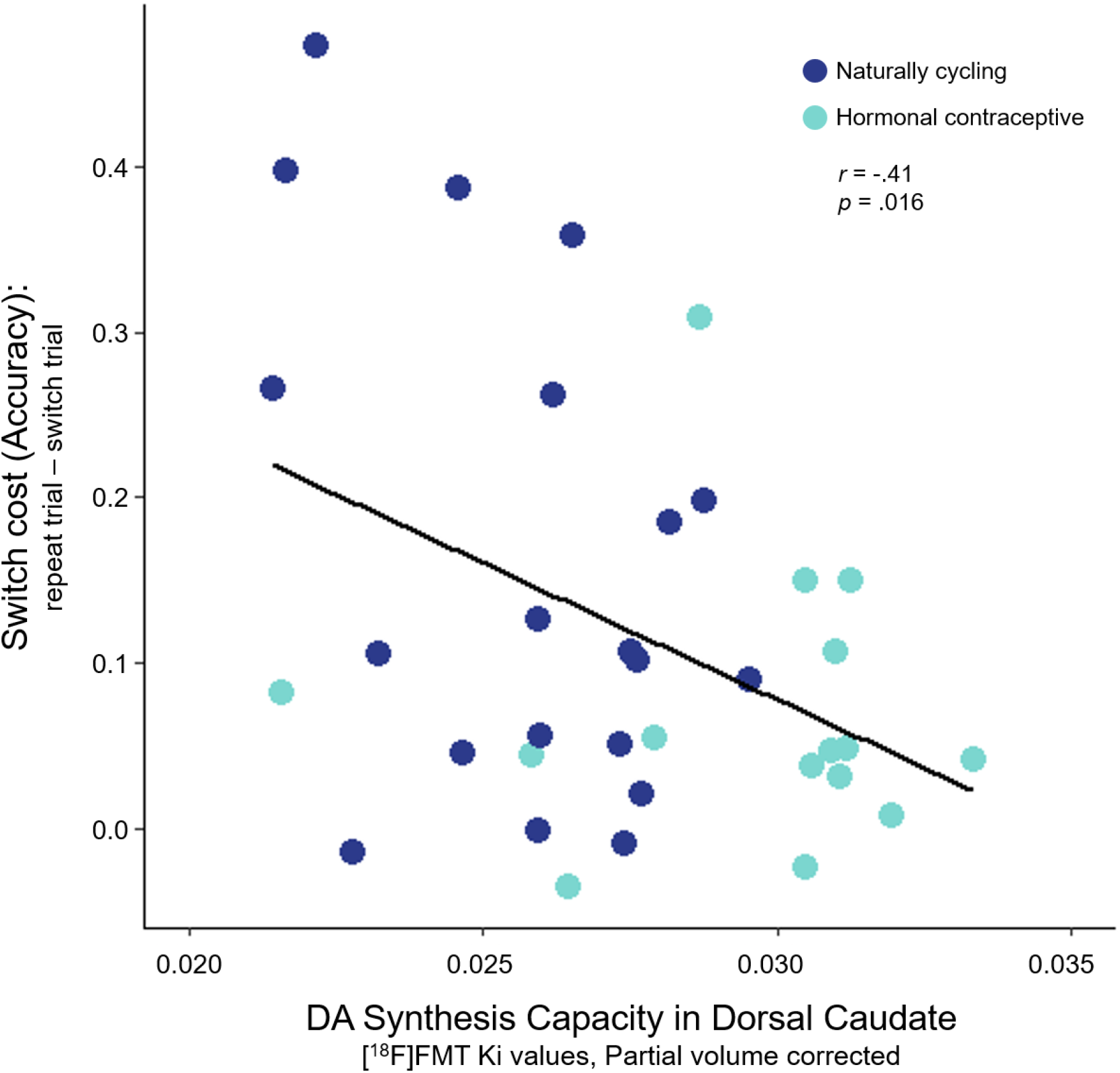
Cognitive flexibility correlates with DA synthesis capacity in dorsal caudate in women. We observed a significant negative correlation between performance on a task switching paradigm and [^18^F]FMT Ki values in dorsal caudate across our female participants (both NC and HC).

#### Men vs Women

We did not observe a difference in switch cost (*t*(46.4) = −0.11, *p* = .91) or distractor cost (*t*(47.9) = −0.47, *p* = .64) between men and women (**Table 3**).

**Table 3.** Performance on task switching paradigm by group

|  | **Distractor Cost** | **Switch Cost** |
| --- | --- | --- |
| **Men** (n= 20) | 0.035 ± 0.040 | 0.125 ± 0.108 |
| **Women** (n = 36) | 0.029 ± 0.051 | 0.121 ± 0.132 |
| **Naturally Cycling**<br>(NC, n = 21) | 0.029 ± 0.054 | 0.160 ± 0.149 |
| **Hormonal Contraception**<br>(HC, n = 15) | 0.030 ± 0.048 | 0.070 ± 0.085 |
| **NC vs HC**<br>cohen's d (Welch's *p*) | n.s. | -0.74 (.03) |
| **Men vs Women** | n.s. | n.s. |
| **Men vs NC vs HC**<br>Kruskal–Wallis *p* | n.s. | n.s. |

#### Men vs Naturally Cycling vs Hormonal Contraceptive Users

We did not observe significant effects of switch cost (*F*(2,52) = 2.428, *p* = .098) or distractor cost (*F*(2,52) = 0.1, *p* = .905) between groups (**Table 3**).

## Discussion

In this study, hormonal contraceptive users exhibited greater dopamine synthesis capacity (as measured by [^18^F]FMT K_i__) and greater cognitive flexibility than naturally cycling participants. No group differences in D2/3 binding potential ([^11^C]raclopride BP) or DA release ([^11^C]raclopride BP PSC) were observed. Though synthesis capacity differed significantly between naturally cycling women and women on hormonal contraceptives, women overall did not differ appreciably from men. This suggests that investigations into the influence of sex hormones on DA neurotransmission may be hampered if limited to comparisons between sexes. Together, these findings lay the groundwork for understanding how global manipulations of the endocrine system, e.g. via hormonal contraceptives, impact dopamine neurotransmission and related cognition.

### DA synthesis capacity differs by hormone status

Though analyses by sex did not reveal differences in DA neurotransmission, when we applied a more nuanced lens to the investigation of hormonal influence on DA function, we found that DA synthesis capacity differed between hormonal contraceptive users and naturally cycling participants, while D2/3 receptor binding potential and stimulated DA release did not differ between groups. These findings are consistent with the preclinical literature. For example, in an ablation-replacement study in ovariectomized rats (11), 17β-estradiol add-back selectively increased striatal DA synthesis but not release, as measured via local superfusion of E2 into the caudate nucleus. Similarly, Algeri et al. (57) observed increased DA synthesis in the striatum and forebrain of intact rats after acute (4 days) and chronic (30 days) oral administration of a synthetic progestin and an estrogen.

Estradiol’s influence on DA synthesis capacity may be mediated by estradiol-induced increases in phosphorylation of tyrosine hydroxylase (TH) (11), the enzyme that converts tyrosine to L-dihydroxyphenylalanine (L-DOPA). Another mechanism of action may be the hormonal regulation of aromatic L-amino acid decarboxylase (AADC) that converts L-DOPA to DA (and is the target of [^18^F]FMT). AADC activity is dependent on pyridoxal phosphate (PLP), or Vitamin B6 (58,59), a nutrient and coenzyme with intermediate concentrations in basal ganglia (60) that is reduced, in some cases to the point of deficiency, in HC users (61–64). If low levels of PLP are associated with reduced AADC activity (65), we would expect HC users to exhibit *reduced* [^18^F]FMT binding relative to naturally cycling women. We observed the opposite pattern. Without information regarding vitamin B6 status for participants, the relationship between PLP and [^18^F]FMT binding remains untested.

The selectivity of our findings to differences in AADC activity (as measured with [^18^F]FMT) and not DA release or D2/3 receptor binding (both measured with [^11^C]raclopride) also suggests the possibility that other catecholamine systems may be impacted. AADC is a critical enzyme in the formation of catecholamines in general, including serotonin (60). In rodent studies, chronic treatment with oral hormonal contraceptives increases brain levels of tryptophan and serotonin (66–67, reviewed in 68). Future investigations should clarify whether global sex steroid hormone manipulations alter DA synthesis capacity specifically, or the catecholamine system generally.

While [^18^F]FMT is a straightforward measure of AADC enzyme activity, which should directly reflect DA synthesis, RAC is a more complex signal. RAC combines several measures, including the binding potential or number of D2/3 receptors (Bavail), and the dissociation constant or how probable the ligand–receptor complex is to dissociate (K_D_). One limitation of our study is that naturally cycling participants were not staged according to menstrual cycle phase. DA release and DA-D2 receptor availability vary across the estrus (3,12,13) and menstrual cycles (10), though see (29,36). In ovariectomized rodents, 17β-estradiol administration augments striatal D2 receptor density (B_avail_), but does not influence binding affinity (1/K_D_) (reviewed in (69)). Thus, it is possible that differences in DA release and baseline binding potential between HC users and unstaged NC women exist, but were obscured in our sample. However, data from Smith and colleagues (37) suggest this is unlikely. In their study, DA release (as measured via [^18^F]fallypride paired with D-amphetamine) did not differ between women using hormonal contraception and naturally cycling women staged within the first 10 days of their menstrual cycle.

Another consideration is that FMT signal increases over the adult lifespan. Braskie et al. (70) observed greater striatal FMT K_i_ values in older participants (mean age = 67) relative to younger participants (mean age = 23). In young adults, higher FMT K_i_ values in caudate are associated with increased working memory capacity (7). In contrast, in older adults greater striatal FMT signal may reflect potential compensation for deficits elsewhere in the DA system (e.g. prefrontal cortex). In a recent study of DA synthesis and working memory capacity in cognitively normal older adults, we (71) observed that adults with the highest FMT K_i_ values also display the greatest atrophy in posterior parietal cortex, raising the possibility of a compensatory response with aging. In the present study of younger adults, HC users were slightly older than NC participants (2 years on average), but the age range of our sample was limited (18–28 years) and results remained significant after controlling for age. Thus, it is unlikely that the group differences we observed are attributable to general effects of aging. Further, our results do not support the idea that higher FMT K_i_ values reflect suboptimal DA functioning, given that HC users had higher FMT K_i_ values and greater cognitive flexibility. Higher FMT K_i_ values in young adults have consistently been associated with better cognitive flexibility (39,72) as well as with working memory capacity (7).

### Consistent effects across hormonal contraceptive regimens

The women in our HC group were on different forms of hormonal contraception, including the combined oral contraceptive pill, vaginal ring, subdermal implant, injection, and hormonal IUD. Exploratory analyses suggest that the relationship between HC use and potentiated DA synthesis capacity is independent of route of administration (**Supplemental Figure 1**). Hormonal contraception (HC) can alter endogenous hormone concentrations to varying extents depending on the formulation and method of delivery. Oral contraceptives and the depot medroxyprogesterone injection exert powerful and sustained suppression of endogenous sex hormone production (73–75), while hormonal IUDs and implants generally exert less pronounced suppression of endogenous hormone levels (75–79). It is possible that the impact of HC on DA occurs via altering endogenous hormone levels, but is likely not solely attributable to endogenous hormone suppression, per se.

The synthetic hormones introduced by the HC regimen, not the alteration of endogenous hormones alone, may be driving changes within the DA system. In one of the few studies of synthetic hormones’ effects on striatal DA, Jori & Dolfini (80) report decreased striatal DA levels in intact female rats after acute and chronic oral administration of steroid contraceptive drug combinations (mestranol with either lynestrenol, norethindrone or norethynodrel). While we did not observe differences in DA receptor binding potential or release, the direction of the effect on DA synthesis capacity that we observed was similar between users of oral contraception (“the pill”, which is primarily a combination of ethinyl estradiol and progestin) and users of other forms of hormonal contraception (including implants, injection, and hormonal IUDs) that primarily contain progestin. This suggests that the progestin component, alone or in concert with endogenous or exogenous estrogen, could be influencing the observed effects. A general consensus from animal and human research is that endogenous estradiol augments DA function (reviewed in (2)), while the influence of progesterone has not been fully characterized (24). Still, progesterone receptor expression in embryonic DA neurons suggests a potentially powerful role of progesterone in modulating DA signaling. In a study of mouse embryonic stem cells, Diaz and colleagues (81) studied the expression of steroid hormone receptors in differentiated DA neurons. They report that 92% of DA neurons expressed progesterone receptors and only 19% of these neurons co-expressed tyrosine hydroxylase and ER-α. Other studies report effects of progesterone, independent of estrogens, on DA release (14,82). Future investigations delineating the influence of synthetic progestins alone and in combination with ethinyl estradiol on DA-ergic function will provide mechanistic insight into the results reported here.

### Hormonal modulation of dorsal caudate vs striatum broadly

We observed a significant difference in DA synthesis capacity between HC and NC groups across the striatum, and post-hoc tests revealed the strongest effect to be in dorsal caudate (**Figure 1**). Thus, it remains unclear whether the effects of hormonal contraception are specific to dorsal caudate, or broadly alter striatal DA synthesis capacity. In a case study of oral contraceptive–induced hemichorea using ^18^FDG-PET, investigators observed striatal hypermetabolism, with increased glucose metabolism in the body of the left caudate nucleus (contralateral to the dyskinesia) (83), suggesting certain caudate-specific effects of oral contraception.

### Individual differences in dopamine synthesis capacity are tied to cognitive flexibility

Hormonal contraceptive users differed from naturally cycling women on switch cost but not on distractor cost in this task-switching paradigm, suggesting a specific effect on cognitive flexibility. This reduced switch cost (i.e., greater cognitive flexibility) in hormone users is consistent with our observation of greater striatal DA synthesis capacity in hormone users relative to naturally cycling women. Previous studies have reported an association between task switching performance and DA synthesis capacity, specifically in the dorsal caudate (39,72,84). Our results suggest an influence of hormonal contraceptive use on the corticostriatal circuitry underlying executive functioning. Future studies should consider whether other measures of executive functioning are influenced, and, by extension, whether dopaminergic medications used to treat disorders of executive function (e.g. ADHD) exert unique effects with or without concomitant use of hormonal contraception.

### Strengths and Limitations

Together, this study provided a unique opportunity to examine differences in basal dopamine receptor occupancy, stimulated dopamine release, and dopamine synthesis capacity in a single cohort, based on women’s hormonal contraceptive status. However, a number of limitations should be considered. First, naturally cycling participants were not staged according to menstrual cycle phase, and as a result we may not have had sensitivity to detect differences in DA signaling between contraceptive users and women at different phases of the menstrual cycle (as opposed to naturally cycling women generally). Second, the route of administration and formulation of the hormonal contraceptive regimen varied (e.g. patch, pill, IUD, implant). Detailed information on participants age of initiation and duration of hormone use would enhance our understanding of the time course with which hormonal contraceptives impacts the DA system.

## Conclusions

This PET imaging study revealed differences in dopamine synthesis capacity between hormonal contraceptive users and naturally cycling women. Measures of DA binding potential and stimulated DA release were similar between groups. Hormonal contraception (in the form of pill, shot, implant, ring or IUD) is used by ~400 million women worldwide (38), yet few studies have examined whether hormonal manipulations impact basic properties of the dopamine system. Findings from this study begin to address this critical gap in women’s health. Moving forward, it is important to consider hormone use as a factor in studies of DA function. More broadly, our findings motivate consideration of the clinical implications of concomitant use of commonly used DA-based medications and hormonal contraceptives.

## End Notes

## Acknowledgements

This work was supported by NIH AG044292 (W.J.), the Daryl and Marguerite Errett Discovery Award (C.M.T), and a NARSAD Young Investigator Grant from the Brain & Behavior Research Foundation (E.G.J).

## Author contributions

The overall project was conceived by M.T.D. and W.J.J., with study aims conceived by C.M.T. and E.G.J.; W.J.J., D.J.F., R.L.W. and A.S.B. and performed the experiments; data analysis was conducted by C.M.T. and D.J.F; C.M.T. and E.G.J. wrote the manuscript; M.T.D., A.S.B., D.J.F., W.J.J., R.L.W., C.M.T, and E.G.J. edited the manuscript.

## Conflict of interest

The authors declare no competing financial interests.

**SI.** Supplementary information is available at MP’s website

