## Supplementary Information for "Striatal dopamine synthesis and cognitive flexibility differ between hormonal contraceptive users and non-users"

**Supplemental Table 1.** DA-D2 binding potential and DA release by group and striatal region of interest

**[$^{11}$C]Raclopride BP Values**

| | Dorsal Caudate | Dorsal Putamen | Ventral Striatum |
|---|---|---|---|
| **Male** | 6.2048 ± .9022 | 7.2033 ± .7939 | 4.0416 ± .4809 |
| **Female (combined)** | 6.0164 ± .1005 | 7.3925 ± .0874 | 3.9975 ± .5846 |
| ***Naturally cycling*** | 6.0122 ± .1182 | 7.4058 ± .7148 | 3.8966 ± .5602 |
| ***Hormonal Contraceptive*** | 6.0231 ± .4048 | 7.3711 ± .5449 | 4.1604 ± .6083 |

**[$^{11}$C]Raclopride BP PSC Values**

| | Dorsal Caudate | Dorsal Putamen | Ventral Striatum |
|---|---|---|---|
| **Male** | .0881 ± .0610 | .1061 ± .0828 | .0664 ± .0585 |
| **Female (overall)** | .0578 ± .0593 | .1142 ± .0755 | .0818 ± .0822 |
| ***Naturally cycling*** | .0586 ± .0553 | .1159 ± .0779 | .0798 ± .0960 |
| ***Hormonal Contraceptive*** | .0565 ± .0676 | .1114 ± .0746 | .0851 ± .0563 |

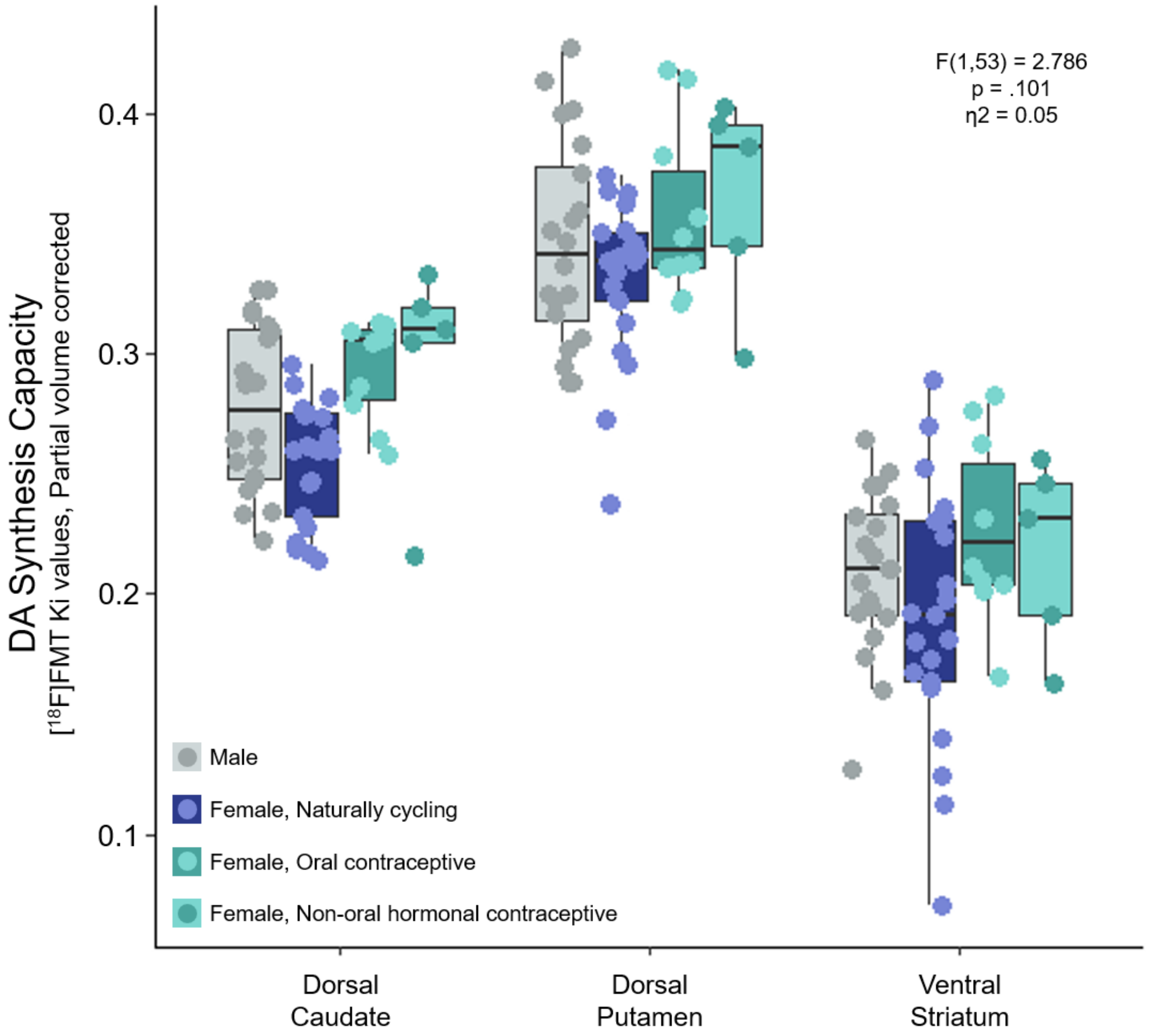


**Supplemental Figure 1. Effects of hormonal contraceptive use on DA synthesis capacity similar with oral and non-oral administrations.** [$^{18}$F]FMT Ki values in males, naturally cycling females, oral contraceptive users (pill; n = 10), and hormonal contraceptive users (i.e. ring, implant, injection, IUD; n = 5), by striatal region of interest. Note that hormone users show elevated DA synthesis, despite the route of administration.
